## supplemental file for "Optimizing Oyster Breeding with Machine Learning and Big Data for Superior Quality"

### Supplementary File 2: Machine Learning Model Parameters and Workflow

#### 1. Machine Learning Algorithms Used

The following machine learning models were employed for predicting oyster breeding traits:

- Random Forest (RF)
- Support Vector Machine (SVM)
- Convolutional Neural Networks (CNNs)

These models were selected due to their robustness in handling complex, high-dimensional biological datasets.

#### 2. Hyperparameters Used

Random Forest:

- Number of estimators: 500
- Maximum depth: 20

Support Vector Machine:

- Kernel: RBF
- Regularization parameter (C): 1.0
- Gamma: Scale

Convolutional Neural Networks (CNNs):

- 3 Convolutional layers with ReLU activation
- 2 Dense layers
- Learning rate: 0.001
- Batch size: 32
- Epochs: 100

#### 3. Data Preprocessing Steps

- Normalization of continuous variables (e.g., shell length, growth rate)
- One-hot encoding of categorical genetic variants
- Train-test split with an 80-20% ratio
- Handling missing values through median imputation

### Supplementary File 2: Machine Learning Model Parameters and Workflow

#### 4. Feature Selection Method

Recursive Feature Elimination (RFE) was applied to identify the most significant features impacting survival probability and meat quality. Feature importance from Random Forest was also used to rank genetic markers.

#### 5. Model Evaluation Metrics

Model performance was evaluated using the following metrics:

- Accuracy
- Precision
- Recall
- F1-Score
- Precision-Recall (PR) curves

Cross-validation with k=5 folds ensured the robustness of the results.

#### 6. Workflow Overview

The machine learning workflow followed these steps:

1. Data Collection: Genomics, transcriptomics, proteomics, and environmental data.
2. Data Preprocessing: Cleaning, normalization, and encoding.
3. Feature Selection: RFE and feature importance analysis.
4. Model Training: RF, SVM, CNN models trained on historical oyster data.
5. Model Evaluation: Metrics calculated and PR curves plotted.
6. Prediction & Validation: Model outputs compared with experimental breeding results.
